# A subpopulation of astrocyte progenitors defined by Sonic hedgehog signaling

**DOI:** 10.1101/2020.06.17.157065

**Authors:** Ellen Gingrich, Kendra Case, A. Denise R. Garcia

## Abstract

The molecular signaling pathway, Sonic hedgehog (Shh), is critical for the proper development of the central nervous system. The requirement for Shh signaling in neuronal and oligodendrocyte development in the developing embryo are well established. Here, we show that Shh signaling also operates in a subpopulation of progenitor cells that generate cortical astrocytes. In the neonatal brain, cells expressing the Shh target gene, *Gli1*, are found in the subventricular zone (SVZ), a germinal zone harboring astrocyte progenitor cells. Using a genetic inducible fate mapping strategy, we show that these cells give rise to half of the cortical astrocyte population, suggesting that the cortex harbors astrocytes from different lineages. Shh activity in SVZ progenitor cells is transient but recurs in a subpopulation of mature astrocytes localized in layers IV and V in a manner independent of their lineage. These data identify a novel role for Shh signaling in cortical astrocyte development and support a growing body of evidence pointing to astrocyte heterogeneity.

## INTRODUCTION

Astrocytes encompass a diverse population of cells that possess a broad array of functional properties that are essential for nervous system function. The developmental processes that confer their functional characteristics are not well understood. While a large body of work has delineated molecular mechanisms underlying astrocyte specification during embryonic development, the processes surrounding postnatal astrocyte development, the time during which most astrocytes are generated, are considerably less well defined.

The Sonic hedgehog (Shh) signaling pathway is best characterized during embryonic neurodevelopment, where it exerts powerful influence over a broad array of neurodevelopmental processes, including patterning and morphogenesis, axon pathfinding, and cell type specification of ventral motor neurons and oligodendrocytes^1^. However, in the adult brain canonical Shh activity is found predominantly in astrocytes^2,3^. Little is known about the role of Shh signaling in astrocyte development. In the developing optic nerve, SHH derived from retinal ganglion cells promotes proliferation of astrocyte precursors^4^. In contrast, SHH in the early embryonic spinal cord limits the specification of astrocyte progenitor cells^5^. Whether Shh signaling plays a role in cortical astrocyte development, is not known.

Astrocyte production occurs primarily during the first two weeks of postnatal development^6^. They are derived from radial glia and progenitor cells in the ventricular zone (VZ) and subventricular zone (SVZ)^7-11^, respectively, as well as from local proliferation of differentiated astrocytes^12^. The dorsal region of the SVZ harbors progenitor cells that generate both oligodendrocytes and astrocytes^8,13^. The postnatal ventricular and subventricular zones (V-SVZ) harbors a population of progenitor cells that express *Gli1*, a transcriptional target of Shh signaling^14,15^. Gli1-expressing progenitors generate a substantial population of oligodendrocytes that populate the overlying white matter^13^. However, in the mature cortex, a discrete subpopulation of astrocytes express *Gli1*, reflecting high levels of Shh activity^3^. Astrocyte transduction of SHH occurs predominantly in cells residing in layers IV and V^16^, consistent with the localization of SHH in layer V pyramidal neurons^3,17^. The precise relationship between Gli1 progenitors in the postnatal SVZ and transduction of SHH in mature astrocytes is not known. One possibility is that superficial and deep layer astrocytes are derived from distinct progenitor pools, with Gli1 precursors exclusively generating deep layer astrocytes. Indeed, large scale transcriptomic analysis of the postnatal cortex shows that astrocytes in deep cortical layers exhibit specific gene expression profiles that distinguish them from those in superficial layers^18^. Alternatively, Shh activity in mature astrocytes may occur independently of their lineage, suggesting that Shh signaling in progenitor cells is uncoupled from its activity in mature cells.

In this study, we performed fate mapping of progenitor cells expressing *Gli1* in the postnatal brain to determine whether these cells generate cortical astrocytes and mapped their distribution across cortical layers. We found that a subpopulation of astrocyte progenitor cells in the SVZ express *Gli1*. Astrocytes within the Gli1 lineage contribute half of the total cortical astrocyte population and are distributed more broadly than those experiencing active Shh signaling in the mature brain. Furthermore, our data show that lineage does not predict Shh activity in mature astrocytes, suggesting that activity of the pathway in progenitor cells is independent from that in mature astrocytes. This may reflect reutilization of the Shh signaling pathway in cells in a manner that is uncoupled from its roles in their development. Taken together, these data demonstrate that Shh signaling identifies a subpopulation of glial progenitor cells that contribute a substantial proportion of protoplasmic astrocytes to the cortex and may have implications for identifying functional heterogeneity of astrocytes. Moreover, these data add to a growing repertoire of cell type-dependent activities orchestrated by Shh signaling.

## RESULTS

### Astrocytes in the Gli1 lineage are broadly distributed in the cortex

We first examined the expression of Gli1 in the early postnatal mouse cortex in *Gli1*^*nlacZ/+*^ mice carrying a nuclear lacZ in the *Gli1* locus^19^. At postnatal day 0 (P0), there was a pronounced population of *Gli1*-expressing cells in the ventral subventricular zone (SVZ), corresponding to the population of SVZ precursors that generate deep granule neurons and periglomerular cells in the olfactory bulb^15,20,21^; (**Figure 1**). *Gli1*-expressing cells were also observed in the dorsolateral corner of the SVZ (dlSVZ), within the postnatal germinal zone of cortical astrocytes^8^, and in a region inferior to the white matter overlying the ventricles (**Figure 1**). Few cells were observed in the cortex at this age (**Figure 1**). We quantified and mapped the distribution of βGal-labeled cells within the dlSVZ and cortex and found that the cortex harbored relatively few *Gli1*-expressing cells at P0 compared to the dlSVZ (**Figure 1**).

**Figure 1.**
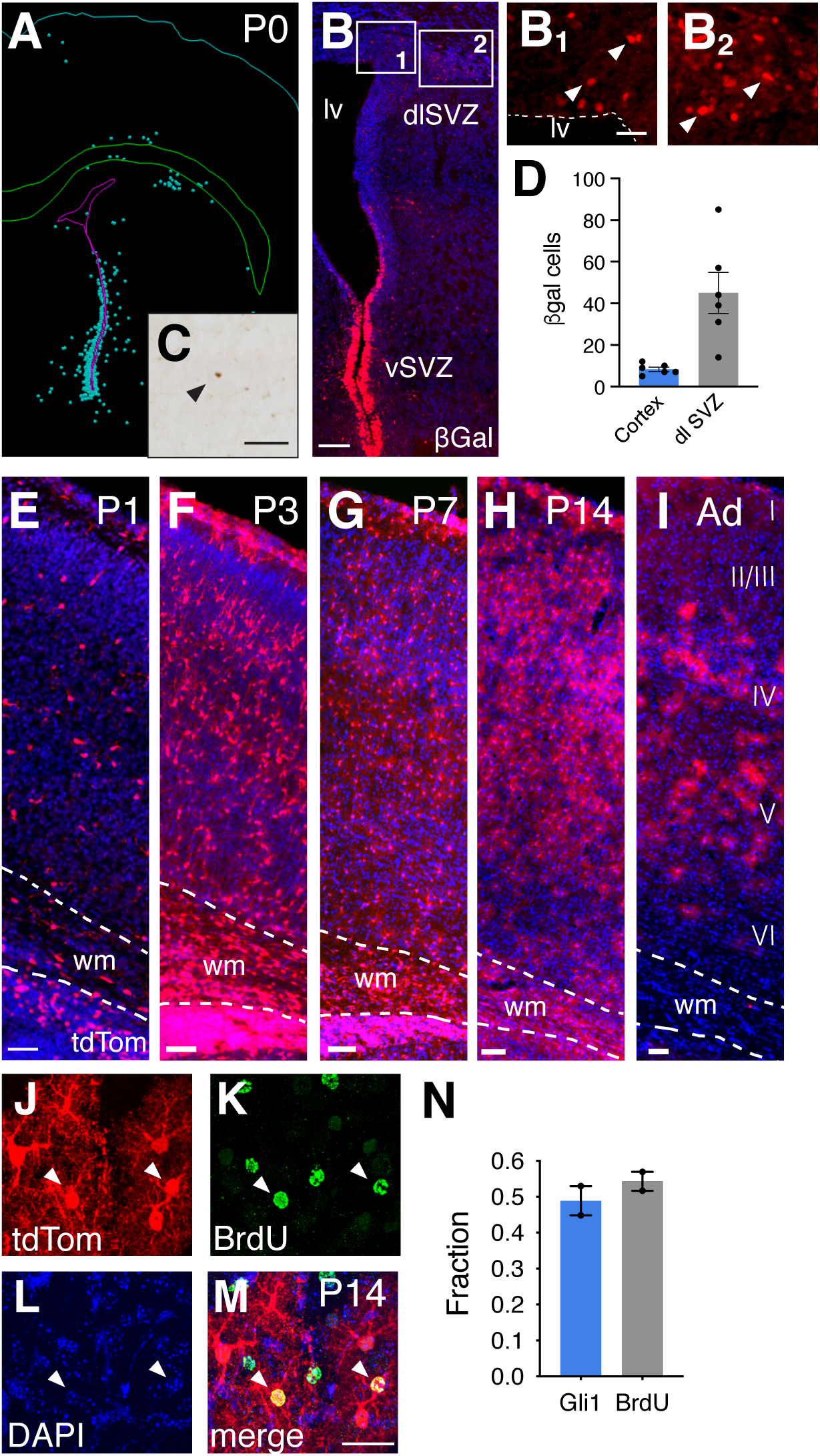
The population of Gli1 cells expands during postnatal development. **(A-D)** The distribution of βGal-labeled cells in *Gli1*^*nlacZ/+*^ tissues at P0. (A) Neurolucida tracing from brightfield immunostained sections. Each dot represents a single cell. (B-C) Immunoflurescent (B) and brightfield (C) staining of βGal at P0 in the SVZ (B) and cortex (C). Insets in B shown in B_1_ and B_2_. Scale bars, B, 100μm, B_1_-B_2_,C, 50μm. lv, lateral ventricle, vSVZ, ventral SVZ, dlSVZ, dorsolateral SVZ. **(D)** Quantification of the number of βGal-labeled cells in the cortex and dlSVZ at P0. Bars represent mean ± SEM, data points represent individual animals. **(E-H)** TdTom (red) expression in the cortex of *Gli1*^*CreER/+*^;Ai14 mice that received tamoxifen at P0 and analyzed at P1 (E), P3 (F), P7 (G), and P14 (H) shows a much broader distribution of marked cells than that observed in adulthood. **(I)** Tamoxifen administration in an adult and analyzed 2 weeks later shows marked cells are found predominantly in layers IV and V. Counterstaining with DAPI (blue), wm, white matter. Scale bar, 50 μm. **(J-M)** Immunofluorescent staining for BrdU in the cortex of a mouse at P14 that received tamoxifen at P0. Single channel (J-K) and merged (M) max projection images from confocal stacks in the cortex showing colocalization of marked cells (red, J) and BrdU (green, K). Counterstaining with DAPI (blue, L). Arrowheads, double labeled cells. Scale bar, 25 μm. (**N**) The fraction of marked cells (Gli1) and dividing cells (BrdU) that are double labeled in mice marked at P0 and analyzd at P14 (*n = 919* Gli1 and *n = 820* BrdU cells). Bars represent mean ± SEM, data points represent individual animals.

To investigate whether *Gli1*-expressing precursors observed at P0 generate astrocytes and/or oligodendrocytes, we performed fate mapping in *Gli1*^*CreER/+*^ mice^14^ carrying the Ai14 tdTomato (tdTom) reporter^22^ (*Gli1*^*CreER/+*^;Ai14). Cre-mediated recombination promotes expression of tdTom that is both permanent and heritable. We marked *Gli1*-expressing precursors by administering tamoxifen to *Gli1*^*CreER/+*^;Ai14 mice at P0 and analyzed the distribution of tdTom at various postnatal ages. One day after tamoxifen (P1), there was a substantial population of marked cells in the cortex (**Figure 1**). At P3, there was a dramatic expansion of marked cells (**Figure 1**), suggesting extensive proliferation between P1 and P3. We also observed many residual radial glial fibers and cells with transitional morphologies, consistent with radial glia undergoing transformation into multipolar astrocytes (**Supplemental Fig. 1**). These cells co-express vimentin, a marker of radial glia (**Supplemental Fig. 1**). There was a further expansion observed at P7 (**Figure 1**), suggesting that *Gli1*-expressing cells marked at P0 correspond to actively dividing glial progenitor cells. There was no further expansion in the number of marked cells between P7 and P14, suggesting the cessation of proliferation of cells marked at birth. To confirm the observed expansion was due to proliferation, we administered BrdU to mice at P0 approximately 12 hours after tamoxifen, to ensure sufficient Cre-mediated recombination prior to incorporation of BrdU. We analyzed tissues at P14 and found that 49% of marked cells in the cortex are co-labeled with BrdU (*-* **Figure 1**), indicating that Gli1 cells marked at P0 are dividing. Conversely, 54% of BrdU labeled cells co-expressed tdTom, suggesting that only a fraction of the proliferating glial precursor cell population residing in the cortex at P0 express *Gli1* (**Figure 1**). Notably, the distribution of marked cells at these postnatal ages was remarkably distinct from that observed in the adult. Whereas tamoxifen administered to adult mice (>P90) produces a distinctive pattern of marked cells found predominantly in layers IV and V^16^, marking *Gli1*-expressing cells at P0 produces a broad distribution of marked cells in the cortex with no laminar specificity (**Figure 1**). These data show that cells within the Gli1 lineage divide rapidly during the first week after birth, and occupy a much broader laminar distribution than that observed in mature astrocytes.

The postnatal ventricular-subventricular zone (V-SVZ) harbors a dorsal domain of *Gli1* expressing radial glia that generates oligodendrocytes which populate the white matter^13^. Our results indicate that cells marked at P0 are not restricted to the white matter as we observed a substantial population of marked cells in the cortex as early as 1 day after tamoxifen. To identify the cell types derived from *Gli1*-expressing precursors, we performed colocalization analysis of marked cells in the cortex with glial cell-type specific markers at P14. Although a small fraction of marked cells co-expressed the oligodendrocyte-specific marker, CC1 (**Figure 2**), the vast majority of marked cells corresponded to astrocytes (**Figure 2**). We examined the morphologies of marked cells and found that at early time points, marked cells showed simple morphologies, with one or two processes, consistent with an immature phenotype (**Figure 2**). Over time, marked cells developed an increasingly complex morphology, extending multiple processes. By P14, cells exhibited a bushy appearance, with many fine branchlets in addition to several major primary branches, consistent with an astrocyte morphology (**Figure 2**). Taken together, these data indicate that Shh signaling is not restricted to mature astrocytes, but is active in progenitor cells in the neonatal dlSVZ that gives rise to a substantial proportion of cortical astrocytes. Our data further show that astrocytes within the Gli1 lineage occupy all layers of the cortex.

**Figure 2.**
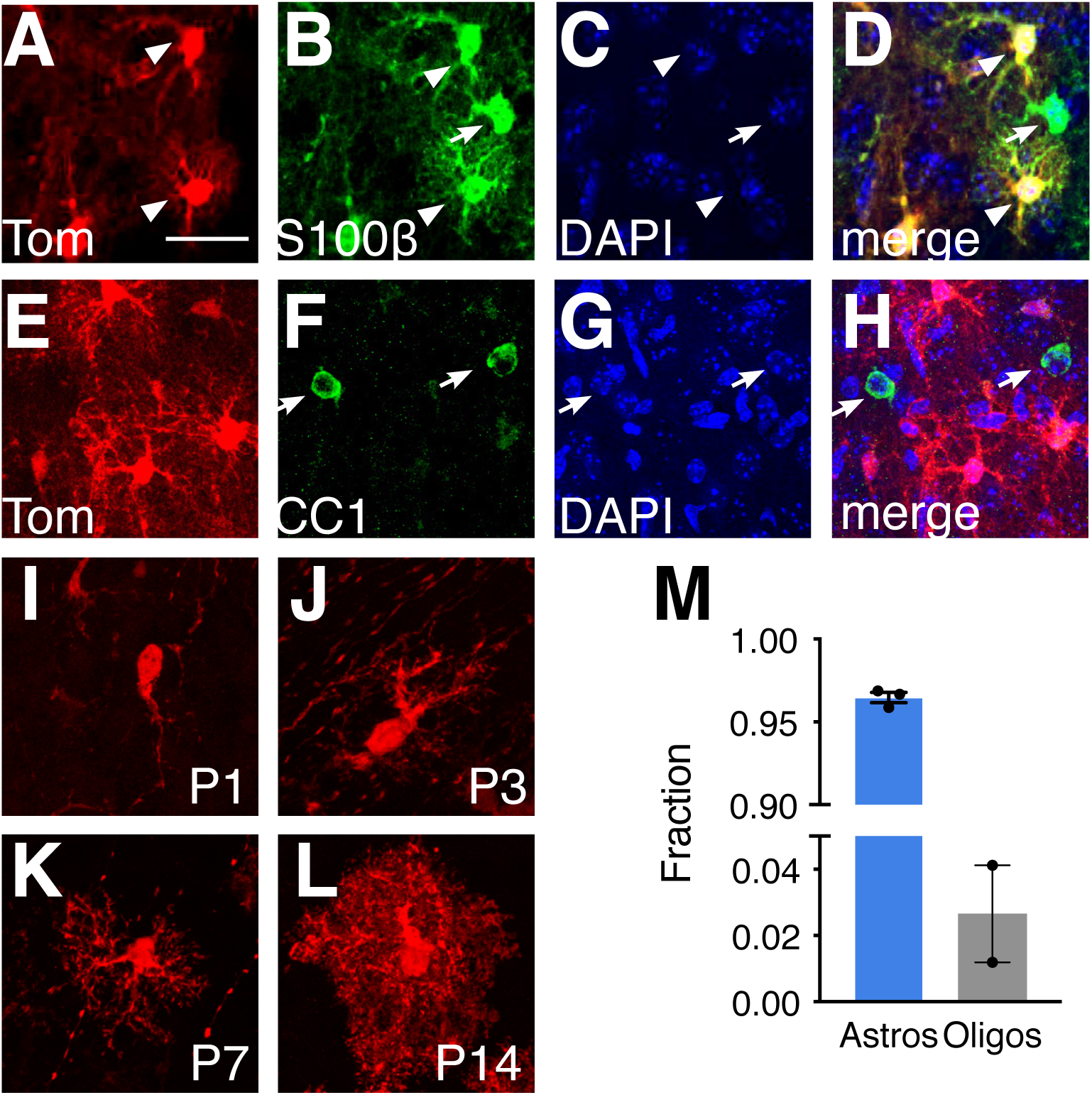
Cells expressing Gli1 at P0 generate cortical astrocytes. **(A-H)** Single channel (A-C, E-G) and merged images (D, H) from confocal images of tdTom (red) and S100β (B, green) or CC1 (F, green) in the cortex at P14, following tamoxifen adminstration at P0. Marked cells show colocalization with S100β, but not with CC1. Counterstaining with DAPI (C, G, blue). Arrowheads, colocalized cells, arrows, single labeled astrocytes or oligodendrocytes not marked with tom. Scale bar, 25 um. **(I-L)** Cells marked at P0, and analyzed at P1 (I), P3 (J), P7 (K), and P14 (L) show an increasingly complex morphology consistent with an astrocyte identity as development proceeds. **(M)** Single cell quantification of the fraction of marked cells at P14 that correspond to astrocytes or oligodendrocytes (*n=783* and *n=632* cells, respectively). Bars represent mean ± SEM, data points represent individual animals.

Whether a Gli1-expressing bi-potent glial progenitor cell generates both cortical astrocytes and white matter oligodendrocytes, or whether these reflect two distinct populations that share Shh activity is not known. Notably, we observed substantial labeling of tdTom in the white matter overlying the ventricles in tissues marked as late as P7 (**Figures 1 and 4**). Further experiments are required to determine whether these cells share a common progenitor or represent two separate populations.

**Figure 3.**
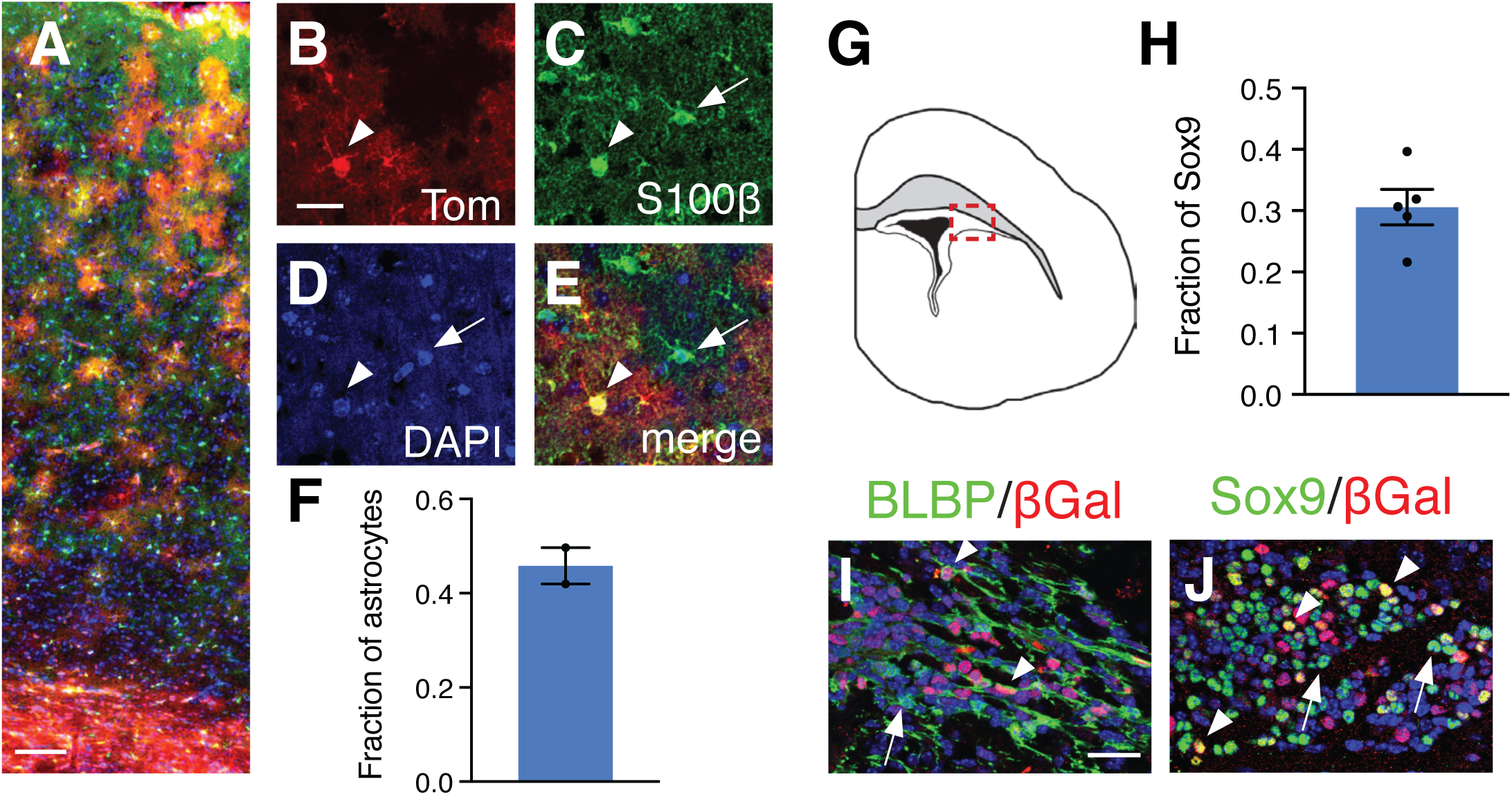
A subpopulation of astrocyte progenitors in the SVZ express Gli1. **(A)** Distribution of tdTom (red) in the cortex of adult (P60) *Gli1*^*CreER/+*^;Ai14 mouse marked at P0. Scale bar, 100 μm. (**B-E**) Single channel, confocal images of tdTom (B, red) and S100β (C, green) identifying marked cells as astrocytes. Counterstained with DAPI (D, blue). Merged image shown in E. Arrowhead, double labeled cell, arrows, single labeled cells. Scale bar, 25 μm (**F**) The fraction of cortical astrocytes in the Gli1 lineage (*n=1371* cells). **(G)** Tracing of a P0 brain section highlighting the dlSVZ (red inset). **(H)** The fraction of Sox9 labeled cells in the dlSVZ co-labeled with βGal at P0 (*n=8358* cells). **(I-J)** Double labeling for βGal (red) and the astrocyte progenitor markers BLBP (I, green) or Sox9 (J, green) in the dlSVZ of *Gli1*^*nlacZ/+*^ mice at P0. Counterstained with DAPI (blue). Scale bar, 25 um. Images taken from red inset shown in G. Bars represent mean ± SEM, data points represent individual animals.

**Figure 4.**
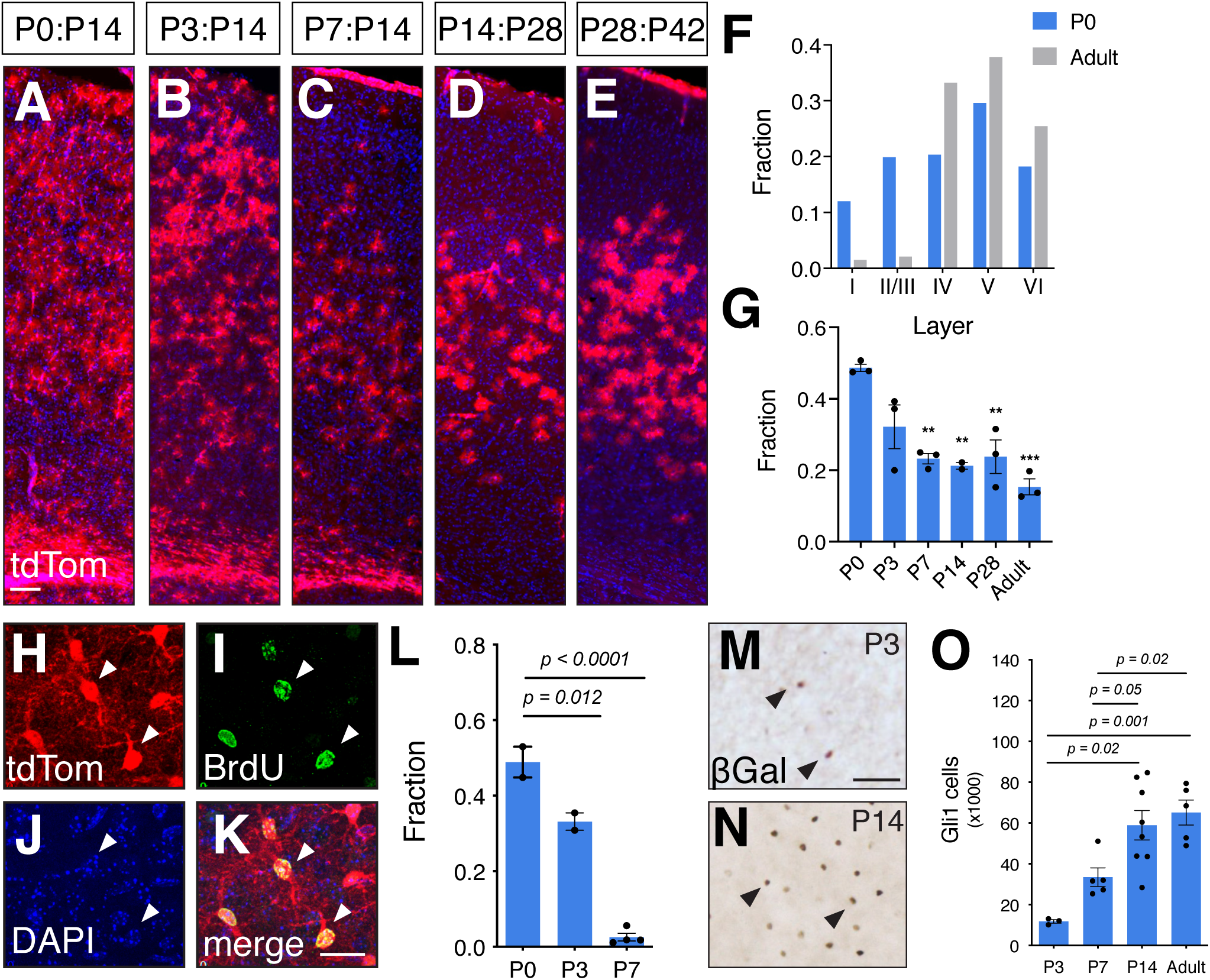
Gli1 cell distribution becomes restricted over postnatal development. **(A-E)** tdTom (red) expression in *Gli1*^*CreER/+*^;Ai14 tissues that received tamoxifen at P0 (A), P3 (B), P7 (C), P14 (D), and P28 (E) and analyzed at ages shown. Counterstaining with DAPI (blue). Scale bar, 100 μm. **(F)** The fraction of marked cells in each layer in mice that received tamoxifen at P0 or in adulthood. **(G)** The fraction of all astrocytes across all layers that are marked at various ages. Bars represent mean ± SEM, data points represent individual animals. ** p < 0.005, ** p < 0.0005 compared to P0. **(H-K)** Single channel, confocal images of tdTom (H, red) and BrdU (I, green) identifying marked cells that are proliferating. Counterstained with DAPI (J, blue). Merged image shown in K. Scale bar, 25 μm. (**L**) The fraction of marked cells colabeled with BrdU at P14 in tissues marked at P0, P3 and P7 (*n=919* cells, *n=720* cells, *n=566* cells, respectively). Bars represent mean ± SEM, data points represent individual animals. (**M-N**) Brightfield immunostaining for βGal in *Gli1*^*nlacZ/+*^ tissues from mice at (M) P3 and (N) P14. Scale bar, 50 μm. **(O)** Estimated total number of Gli1-expressing cells derived from stereological quantification of βGal labeled cells in the cortex of *Gli1*^*nlacZ/+*^ mice at various ages. Bars represent mean ± SEM, data points represent individual animals.

### Gli1 progenitors generate a subpopulation of cortical astrocytes

We next investigated the extent to which astrocytes in the Gli1 lineage contribute to the total astrocyte population in the mature cortex. We administered tamoxifen at P0 and analyzed the fraction of cells labeled with the pan-astrocytic marker, S100β, that co-express tdTom at P60. We found that 46% of astrocytes were co-labeled with tdTom (**Figure 3**), indicating that astrocytes within the Gli1 lineage make up nearly half of the total cortical astrocyte population. This suggests that the cortex harbors a mixed population of astrocytes from different lineages. These data further suggest that *Gli1*-expressing astrocyte progenitor cells comprise a subpopulation of the total pool of astrocyte progenitors. Alternatively, these data could reflect limitations of tamoxifen-dependent Cre-mediated recombination. To rule out the possibility that recombination was inefficient within the *Gli1*-expressing progenitor population, we examined the dlSVZ of *Gli1*^*nlacZ/+*^ mice, enabling us to identify cells actively expressing *Gli1* independent of the requirement for recombination. We identified the pool of astrocyte progenitors at P0 by expression Sox9 and found that 31% co-expressed *Gli1* (**Figure 3**), consistent with the idea that progenitor cells expressing *Gli1* comprise a subpopulation of astrocyte progenitors in the P0 dlSVZ. This was confirmed with a second astrocyte progenitor marker, BLBP, in which we found that many BLBP labeled cells were Gli1 negative (**Figure 3**). Taken together, these data suggest that the astrocyte progenitor pool in the dlSVZ at P0 is comprised of a molecularly distinct subpopulation that can be defined by Shh signaling.

### Shh signaling in the cortex is upregulated during postnatal development

Our fate mapping studies showed a differential distribution of cells marked in the neonatal compared to the adult brain. Most notably, genetic marking at P0 produces many labeled cells localized to superficial layers of the cortex, in contrast to adulthood in which marked cells are found predominantly in deep layers (**Figure 4**). We next investigated the age at which the distinctive laminar pattern of Shh signaling in the cortex emerges. We administered tamoxifen over three consecutive days at various ages during postnatal development. Tissues were analyzed two weeks after the initial tamoxifen dose, with the exception of P7 animals which were analyzed at one and three weeks after tamoxifen. There was no difference between these chase periods and the data were pooled for P7. As with P0, the vast majority of marked cells labeled at these postnatal ages were subsequently identified as astrocytes (**Table 1**). Analysis of the fraction of marked cells in each layer shows that cells marked at P0 are distributed throughout all layers, whereas the vast majority of cells marked in the adult brain are found in deep layers (**Figure 4**). At P3, the distribution of marked cells remains broad throughout all cortical layers. By P7, fewer marked cells are found in the upper cortical layers than in tissues marked at younger ages (**Figure 4**). However, by P14 marked cells are found predominantly in deep layers and are largely absent from superficial layers, a pattern that was consistent with that observed at P28 and beyond (**Figure 4**). This was associated with a concomitant reduction in the fraction of astrocytes, identified by S100β, that are marked at these ages (**Figure 4**). At P0, 49% of astrocytes across all layers are marked, but by P3, that proportion declined to 32%, though this was not significant (**Figure 4**). At P7, the fraction of marked astrocytes declined significantly from P0 to 23% This fraction remained steady at P14 and P28 (21% and 24%, respectively; 15%, **Figure 4**). There was a further modest reduction to 15% in adults, though this was not significant (**Figure 4**). Double labeling with BrdU administered 12 hours after tamoxifen shows that the fraction of marked cells that are dividing declines dramatically over the first postnatal week. At P0, the fraction of marked cells double labeled with BrdU was 49%, whereas at P3, that fraction was significantly reduced to 33%. By P7, nearly all marked cells were postmitotic as only 3% of tdTom cells co-labeled with BrdU (**Figure 4**). This suggests that astrocytes within the Gli1 lineage are generated predominantly during the first few days after birth.

**Table 1.** Gli1 cells marked at various ages correspond to astrocyte progenitors or mature astrocytes. Single cell analysis of the identity of marked cells in *Gli1*^*CreER/+*^;Ai14 mice that received tamoxifen at the ages shown in the left column, and analyzed two weeks later. The fraction of tdTom cells identified as astrocytes by colocalization with S100β. The number of individual cells analyzed is in parentheses. *n = 3 - 4* mice for each age.

| Age | Astrocytes |
| --- | --- |
| P0 | 95% (783) |
| P3 | 99% (843) |
| P7 | 99% (578) |
| P14 | 97% (348) |
| P28 | 98% (389) |
| Adult | 99% (191) |

Despite the progressive decline in the fraction of astrocytes that are marked during the first postnatal week, analysis of active Shh signaling the cortex in *Gli1*^*lacZ/+*^ mice shows a progressive increase over the first two postnatal weeks (**Figure 4**). Stereological quantification of the number of βGal labeled cells in the cortex showed a significant increase in the number of cells at P14 compared to P3 (**Figure 4**). There was no difference between P14 and the adult (>P90) cortex (**Figure 4**), suggesting that Shh signaling in the cortex stabilizes by the second postnatal week. Despite the increase in the number of cells exhibiting Shh activity between P7 and adulthood, few cells marked at P7 are proliferating (**Figure 4**), arguing against the possibility that the increase in *Gli1*-expressing cells between P7 and adulthood is due to proliferation of immature progenitor cells. These data suggest that Shh signaling in astrocyte progenitor cells residing in the SVZ is transient and is lost as cells migrate into the cortex and undergo maturation. In parallel, Shh activity in the cortex is low at birth, but increases during postnatal development in a subpopulation of mature, postmitotic astrocytes found mostly in deep layers, coincident with the localization of Shh-expressing neurons in layer V^17,23^. Taken together, these data suggest that Shh signaling in mature cortical astrocytes operates independently of that in SVZ progenitor cells.

### Lineage does not predict Shh activity in mature astrocytes

We next examined whether Shh signaling in mature astrocytes is restricted to those within the Gli1 lineage. We crossed *Gli1*^*CreER/+;*^*Ai14* mice with *Gli1*^*nlacZ/+*^ mice (*Gli1*^*CreER/nlacZ*^;*Ai14*) to generate mice in which we could distinguish between temporally distinct populations of cells expressing *Gli1*. We reasoned that because tamoxifen administered at P0 will indelibly mark progenitor cells expressing *Gli1* and their progeny, whereas βGal-expressing cells would reflect *Gli1* activity at the conclusion of the experiment, this approach would enable us to identify individual cells showing differential *Gli1* activity at two different time points within a single mouse. While this effectively produces a *Gli1* null mouse, *Gli1* is not required for Shh signaling during development and *Gli1* null mice show no developmental or behavioral deficits^19^. We administered tamoxifen to *Gli1*^*CreER/nlacZ*^;*Ai14* mice at P0 and analyzed tissues at P60 for colocalization of the tdTom and βGal reporter proteins. Although cells marked at P0 are found throughout all cortical layers, this analysis was restricted to deeper layers where βGal-expressing cells are mostly found. At P60, a large proportion (67%) of tdTom-labeled cells did not co-express βGal (**Figure 5**), suggesting that in a substantial fraction of astrocytes in the Gli1 lineage, Shh signaling is downregulated sometime after P0. The remaining fraction of tdTom-labeled cells were double labeled with βGal (**Figure 5**), suggesting either persistent Shh activity since birth or re-induction of *Gli1* in the same cell. Interestingly, 40% of βGal-labeled cells did not co-express tdTom (**Figure 5**), suggesting active Shh signaling in mature astrocytes that are not within the Gli1 lineage. Taken together, these data suggest that Shh signaling in differentiated astrocytes reflects recurrence of Shh activity in mature cells that is independent of pathway activity in postnatal progenitor cells residing in the SVZ.

**Figure 5.**
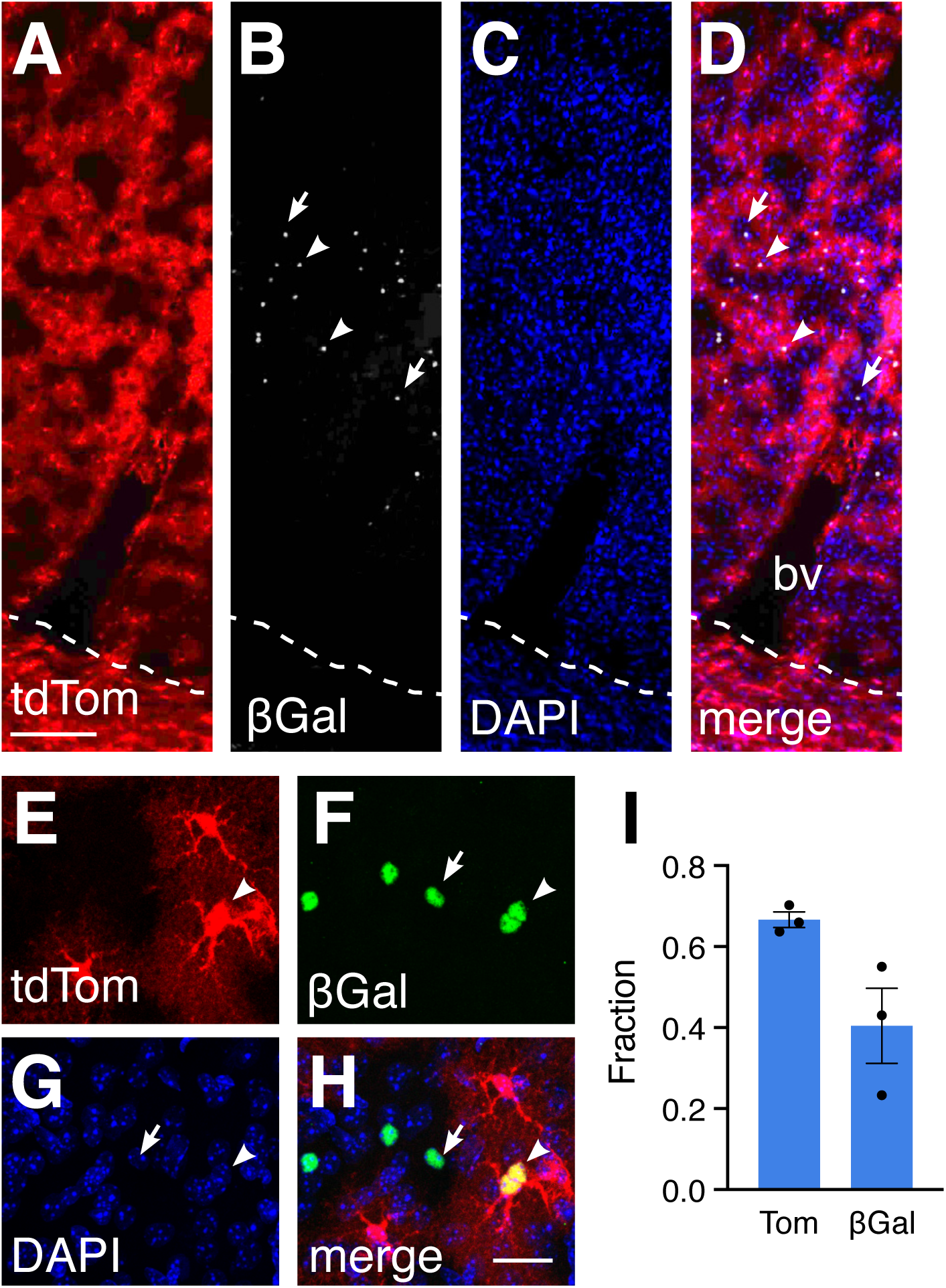
Shh signaling is upregulated in mature astrocytes. **(A-H)** Double labeling for tdTom (A, E, red) and βGal (B, grey, F, green) in the cortex of a *Gli1*^*CreER/nLacZ*^;Ai14 mouse marked at P0 and analyzed at P60. (A-D) Low magnification and (E-H) high magnification confocal images. Counterstained with DAPI (C, G, blue). Dotted line depicts white matter border. Merged images shown in D, H. bv, blood vessel. Arrowheads, colocalized cells, arrows, βGal only cells. **(I)** The fraction of tdTom or βGal labeled cells that are single labeled (*n=1454* and *n=821* cells, respectively). Bars represent mean ± SEM, data points represent individual animals.

## DISCUSSION

Using genetic fate mapping together with a direct reporter of Shh activity, we identified Shh signaling in progenitor cells that generate a substantial proportion of cortical astrocytes. Shh activity is transient and is lost as postnatal development proceeds. Astrocytes within the Gli1 lineage contribute half of all cortical astrocytes, consistent with the observation that Shh is active in a subpopulation of astrocyte progenitors in the neonatal brain. Astrocytes within the Gli1 lineage are distributed broadly across superficial and deep cortical layers, in contrast to the laminar pattern of Shh signaling in the mature cortex. Finally, our data show that lineage does not predict Shh activity in mature astrocytes, suggesting that the role of Shh signaling in mature astrocytes is independent of its role in progenitor cells.

The diversity of mature astrocyte populations has gained increasing recognition, but how such diversity is achieved is not well understood. One mechanism by which the CNS can achieve such diversity is through the production of cells from distinct progenitor cell lineages. In the embryonic spinal cord and forebrain, astrocyte progenitor cells residing in molecularly and anatomically distinct domains produce regionally specified astrocytes that occupy the overlying territory defined by radial glial trajectories^24,25^. Here, we show that Shh signaling operates in a subpopulation of astrocyte progenitor cells residing within the postnatal SVZ.

Our observation that half of all cortical astrocytes are derived from a Gli1 lineage suggests that additional diversity exists among postnatal progenitor cells residing within a single domain. The extent to which lineage confers distinct molecular or functional characteristics in astrocytes is not known. The postnatal V-SVZ harbors neural stem cell populations with distinct positional identities that generate specific subpopulations of olfactory bulb interneurons^26-28^. Whether astrocytes within the Gli1 lineage represent a functionally distinct class of cells is unknown and should be the basis of further studies.

One limitation of our study is the requirement for tamoxifen to trigger Cre-mediated recombination. It is possible that the dose and timing of tamoxifen may be insufficient to label all progenitor cells expressing *Gli1*. However in *Gli1*^*nlacZ/+*^ mice in which expression of the reporter protein is directly regulated by *Gli1* activity, we found that 30% of progenitor cells express Gli1 in these mice. Given this proportion, our observation that half of the astrocyte population in adult is marked is somewhat lower than might be expected. One possibility is that astrocytes undergo developmental elimination. Indeed, astrocytes in the retina are engulfed by microglia during early postnatal development^29^, and the authors further report a developmental decline in cortical astrocytes in the adult. Such elimination of cells during postnatal maturation could underlie our observations and suggests that Gli1-expressing progenitors generate an even larger fraction of cortical astrocytes than we observed in this study. Nevertheless, our observation that only a fraction of astrocyte progenitor cells express *Gli1* at P0 is consistent with the idea that Shh signaling identifies a defined subpopulation of astrocyte progenitor cells. It is possible that the astrocyte progenitor pool at this age may reflect cells in different stages of development. It will be interesting to explore whether timing cooperates with molecular signaling to confer specific functional properties upon astrocytes derived from molecularly and temporally distinct pools of progenitor cells.

Astrocytes are derived from three sources, radial glia, progenitors in the SVZ, and local proliferation of astrocytes in the cortex^7,8,12,30^. Our data show that cells expressing *Gli1* at P0 are found in substantially greater numbers in the dlSVZ, with relatively fewer cells in the cortex. In addition, we observed many radial glial processes as well as cells with transitional unipolar or bipolar morphologies, consistent with the transformation of radial glia into multipolar astrocytes. These data suggest that Shh is operating in radial glial and SVZ progenitors, although we cannot rule out the possibility that Shh signaling is also acting on proliferating cells in the cortex that are expanding the population of Gli1 lineage astrocytes. However if this is the case, then their proliferation capacity appears to be exhausted before P7, as only 3% of marked cells in the cortex are proliferating in animals receiving both tamoxifen and BrdU at this age. This argues against the possibility of local proliferation, and instead suggests that as progenitor cells in the Gli1 lineage migrate away from the SVZ and into the cortex, they lose sensitivity to Shh signaling. This may reflect the availability of SHH to astrocyte progenitors and mature astrocytes and suggests that there may be differential sources of ligand to these cell populations. There are several potential sources of SHH in the neonatal and postnatal brain which include ventral forebrain neurons^15^, cortical neurons^17,31^, epithelial cells^32^, and the CSF^33,34^. Because high levels of SHH are required to stimulate Gli1 expression^19^, our data suggest that *Gli1*-expressing progenitors are migrating away from their neonatal source of SHH. Furthermore, our data suggest that astrocytes within the Gli1 lineage are generated predominantly during the first few days after birth and that Shh activity in astrocyte progenitor cells is transient and declines as astrocytes undergo maturation.

Intriguingly, our fate mapping studies show that while the fraction of astrocytes in the cortex that are marked declines progressively between P0 and P14, the number of cells showing active Shh signaling increases progressively during this time. Shh levels in the postnatal cortex are developmentally regulated and are low at birth but increases progressively until its peak at P14^23^. This suggests that as early postnatal development proceeds, Shh activity in the cortex reflects a recurrence of the pathway in differentiated cells, independently of activity in SVZ progenitor cells. Moreover, lineage does not predict Shh activity in mature astrocytes, as cells that were not marked at P0 nevertheless exhibit Shh activity in adulthood. Consistent with this, nearly all the components of the Shh signaling pathway, including the receptor *Patched* (*Ptc)*, and the transcriptional effectors *Gli2* and *Gli3* are found in a greater number of astrocytes than are actively expressing *Gli1*^*3*^, suggesting that many cells possess the machinery to transduce SHH. This may reflect dynamic activity of the pathway and may be one mechanism by which neurons can recruit neighboring astrocytes depending on local needs.

Shh signaling regulates a diverse repertoire of cellular activities in multiple cell types, including neural stem and progenitor cells, neurons and astrocytes^35^. It is likely that the molecular programs initiated by Shh activity in astrocyte progenitors and mature astrocytes confer distinct functional characteristics that are cell-type dependent. This application of a single molecular pathway in cell type dependent ways reflects the pleiotropic nature of Shh signaling and demonstrates the remarkable capacity of the pathway to be deployed in a broad array of cellular activities.

## MATERIALS AND METHODS

### Animals

The following transgenic mouse lines were used: *Gli1*^*nLacZ/+*,19^, *Gli1*^*CreER/+*,14^, and *R26*^*tdTom/tdTom*^ (Ai14)^36^. Animals were maintained on a 12h light/dark cycle and given access to food and water ad libitum. All experiments were conducted in accordance with the Drexel University Institute for Animal Care and Use Committee. Male and female animals between postnatal day (P) 0 and P60 were used.

### Tamoxifen

Tamoxifen (Sigma, T5648-1G) was diluted to a final concentration of 5mg/ml or 10mg/mL in corn oil. P0 *Gli1*^*CreER/+*^;*Ai14* mice received 50mg/kg tamoxifen by intragastric injection for 1 or 3 days consecutively and tissue was harvested at various indicated time points. All other *Gli1*^*CreER/+*^;*Ai14* mice received 100mg/kg of tamoxifen by intragastric injection (P3), subcutaneous injection (P7), or oral gavage (P14 and above) for 1 or 3 days consecutively, and tissue was harvested at indicated time points. For adult comparisons, *Gli1*^*CreER/+*^;*Ai14* mice greater than 2 months old received 250mg/kg of tamoxifen by oral gavage for 1 or 3 days consecutively, and tissue was harvested 2 weeks later, unless otherwise noted.

### BrdU

BrdU (Sigma, B9285-1G) was dissolved in 0.007N NaOH in sterile saline and administered via intraperitoneal (i.p.) injection. For long term experiments, mice received 50mg/kg in mice ages P0 to P14 or 200mg/kg at P28 and older, at 6 – 24 hours after tamoxifen.

### Perfusion and histology

Animals were given an i.p. injection of a Ketamine/Xylazene/Acepromazine cocktail and transcardially perfused with 10mM PBS followed by 4% paraformaldehyde solution. Brains were dissected and post-fixed in 4% paraformaldehyde for 4-6 hours followed by cryoprotection in 30% sucrose and stored at 4°C for at least 48 hours or until ready for sectioning. For P0 and P3, tissue was embedded in OCT medium after cryoprotection and stored at -20°C until sectioning. This tissue was sectioned by cryostat at 16 - 20μm onto coated slides and stored at -80°C, protected from air and light. All other tissue was sectioned by cryostat at 40μm and stored at 4°C in TBS with 0.05% sodium azide. Immunohistochemistry was performed on every 12^th^ section both for free floating and slide-mounted tissues using the following primary antibodies for fluorescence: rabbit anti-βgal (1:1k/1:10kMP Biomedicals), chicken anti-βgal (1:1k, Abcam), mouse anti-BLBP (1:1k, Abcam), sheep anti-BrdU (1:500, Maine Biotechnology Services), mouse anti-CC1 (1:1k, Calbiochem), rabbit anti-RFP (1:500, MBLI), rabbit anti-S100β (1:1k, DAKO), goat anti-Sox9 (1:1000 R&D), and chicken anti-Vimentin (1:1k, Invitrogen). For BrdU staining, tissue was pre-incubated in 2N HCl for 30 minutes and neutralized with 0.1M TBS before incubation in block and primary antibody. Fluorescent labeling was achieved using species-specific AlexaFluor-tagged secondary antibodies, Alexa Fluor 488, Alexa Fluor 568, or Alexa Fluor 647 (1:1k, Life Technologies), followed by counterstaining with DAPI (1:36k, Life Technologies). For brightfield immunostaining, tissues were quenched in TBS with 0.3% H2O2 and 30% methanol for 30 minutes prior to incubation in block and primary antibody. The following antibodies were used: rabbit anti-βgal (1:40K, MP Biomedicals) and rabbit anti-RFP (1:500, MBLI). For brightfield staining, species-specific biotinylated secondary antibodies (Vector) were used at 1:400 followed by incubation in avidin-biotin complex (ABC, Vector). Visualization was achieved using 3’-3 diaminobezedine (DAB, Vector) as the developing agent. 4% paraformaldehyde post-fix was applied to all slide-mounted tissue for 30 minutes prior to staining.

### Microscopy

Stained sections were examined and imaged in brightfield and fluorescence using an upright microscope (Zeiss) and StereoInvestigator software (MBF Biosciences). Confocal images were obtained on an inverted microscope (Leica) using LAS X software. Single-cell analysis of co-labeling was evaluated on double-stained immunofluorescent tissues by taking confocal sections with a 1um slice distance with 20x, 40x oil or 60x oil objective. For each cortex sampled, multiple z stacks were collected from 3-5 sections within the anterior and posterior boundaries defined by figure 22 and figure 44, respectively from Paxinos and Franklin (2013).

### Cell quantification

Stereological estimates of the total number of cells was performed in tissues stained for brightfield microscopy. The cortex was analyzed in a series of 6-8 sections, sampled every 480µm, and bounded by the midline dorsally and the end of the external capsule ventrally. Anterior and posterior boundaries were defined figure 22 and figure 44, respectively from Paxinos and Franklin^37^. We used a modified optical fractionator and stereological image analysis software (StereoInvestigator, MBF Bioscience) operating a computer-driven stage attached to an upright microscope (Zeiss). The cortical area to be analyzed was outlined at low magnification, and counting frames were selected at random by the image analysis software. Cells were counted using a 40x objective and DIC optics. A target cell count of 300 cells was used to define scan grid and counting frame sizes, with a 2μm guard zone. For all analyses, only cells with a clear and distinct labeled cell body were analyzed. Analysis of cells in the cortex and dlSVZ was performed on tissues stained for brightfield microscopy. Each region was traced in Neurolucida (MBF Bioscience) at 5x and individual βgal-labeled cells were mapped with separate markers in each region at 40x using DIC optics.

### Statistics

All statistical analyses were performed using Prism 8 (GraphPad).

## Acknowledgements

The authors are grateful to Dr. Corey Harwell (Harvard Medical School) for helpful discussions and critical feedback.

---

**Supplemental Figure 1.**
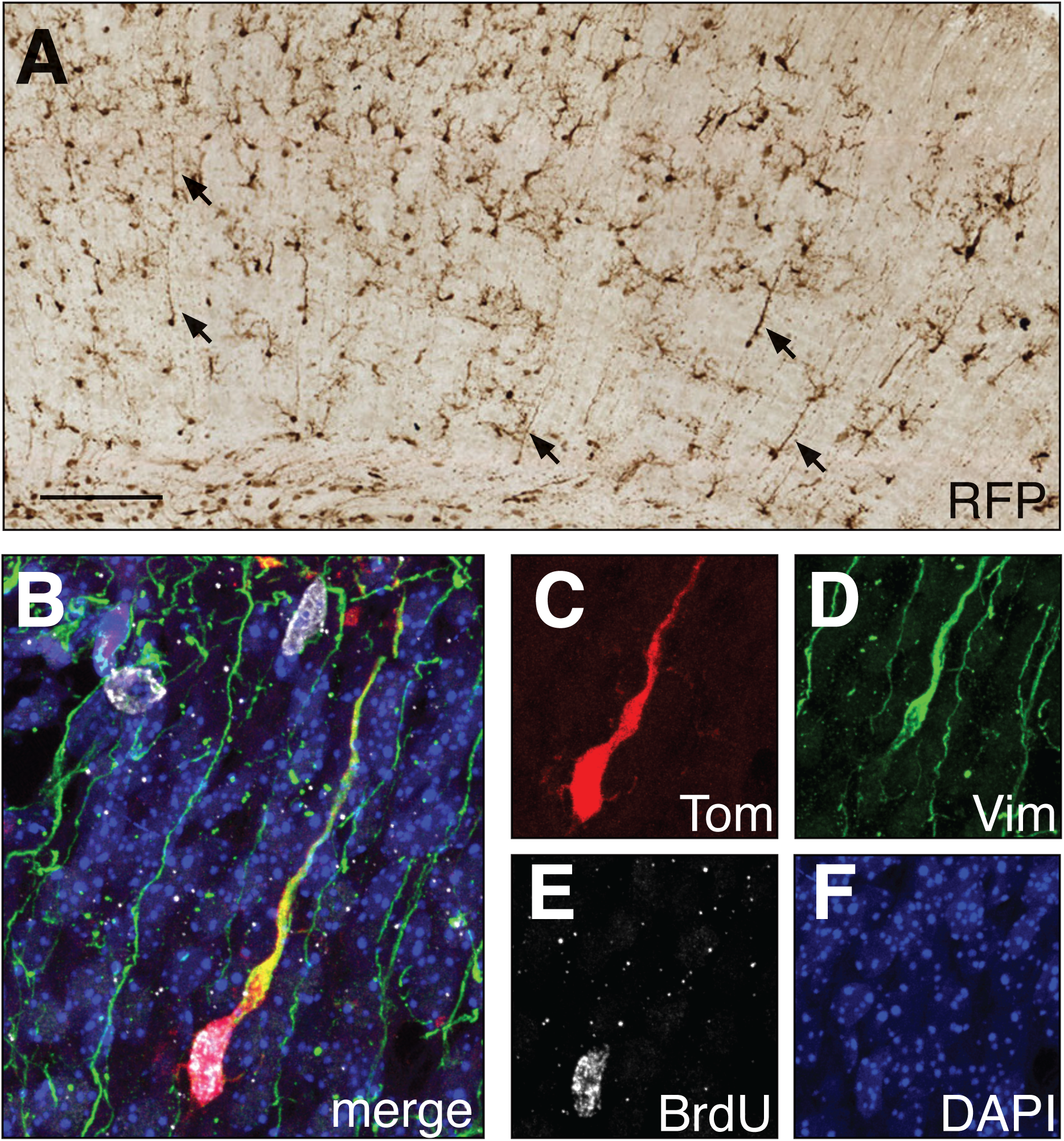
Marked cells show characteristics of transitional radial glia. **(A)** Brightfield immunostaining for RFP in the cortex of a mouse at P3 after receiving tamoxifen at P0 showing many cells with transitional morphologies and the appearance of residual radial glial fibers. Scale bar, 25 μm **(B-F)** Colocalization of tdTom (C, red), vimentin (D, green), and BrdU (E, gray) in the cortex of *Gli1*^*CreER/+*^;Ai14 mice at P3 after tamoxifen at P0. Merged image in (B), single channel images in C-F.

